# Engineered bacteria detect tumor DNA

**DOI:** 10.1101/2021.09.10.459858

**Authors:** Robert M. Cooper, Josephine A. Wright, Jia Q. Ng, Jarrad M. Goyne, Nobumi Suzuki, Young K. Lee, Mari Ichinose, Georgette Radford, Feargal Ryan, Shalni Kumar, Elaine M. Thomas, Laura Vrbanac, Rob Knight, Susan L. Woods, Daniel L. Worthley, Jeff Hasty.

## Abstract

Advances in bacterial engineering have catalysed the development of living cell diagnostics and therapeutics^1–3^, including microbes that respond to gut inflammation^4^, intestinal bleeding^5^, pathogens^6^ and hypoxic tumors^7^. Bacteria can access the entire gastrointestinal tract^8^ to produce outputs measured in stool^4^ or urine^7^. Cellular memory, such as bistable switches^4,9,10^ or genomic rearrangements^11^, allows bacteria to store information over time. However, living biosensors have not yet been engineered to detect specific DNA sequences or mutations from outside the cell. Here, we engineer naturally competent *Acinetobacter baylyi* to detect donor DNA from the genomes of colorectal cancer (CRC) cells, organoids and tumors. We characterize the functionality of the biosensors *in vitro* with co-culture assays and then validate *in vivo* with sensor bacteria delivered to mice harboring colorectal tumors. We observe horizontal gene transfer from the tumor to the sensor bacteria in our mouse model of CRC. The sensor bacteria achieved 100% discrimination between mice with and without CRC. This Cellular Assay of Targeted, CRISPR-discriminated Horizontal gene transfer (CATCH), establishes a framework for biosensing of mutations or organisms within environments that are difficult to sample, among many other potential applications. Furthermore, the platform could be readily expanded to include production and delivery of antibiotic or antineoplastic therapeutic payloads at the detection site.

## Main text

Some bacteria are naturally competent for transformation and can sample extracellular DNA directly from their environment^12^. Natural competence is one mechanism of horizontal gene transfer (HGT), the exchange of genetic material between organisms outside vertical, “parent to offspring” transmission^13^. HGT is common between microbes^13^ and from microbes into animals and plants^14^. Genomic analyses have also found signatures of HGT in the other direction, from eukaryotes to prokaryotes^15^, but the forward engineering of bacteria to detect or respond to human DNA via HGT has not been explored. *Acinetobacter baylyi* is a highly competent and well-studied bacterium^16^ that is largely non-pathogenic in healthy humans^17^ and can colonize the murine gastrointestinal tract^18^. This combination of traits renders *A. baylyi* an ideal candidate for studying engineered detection of colorectal cancer DNA (Fig. 1). Our CATCH strategy delivers bacterial biosensors to the gastrointestinal tract, where they sample and genomically integrate target tumor DNA. To demonstrate the concept, we use the biosensor to detect engineered tumor cells. We then develop genetic circuits to detect natural, non-engineered tumor DNA sequences, discriminating oncogenic mutations at the single base level. Since the target sequence and output gene are modular, our approach can be generalized to detect arbitrary DNA sequences and respond in a programmable manner, in a range of future contexts.

**Figure 1.**
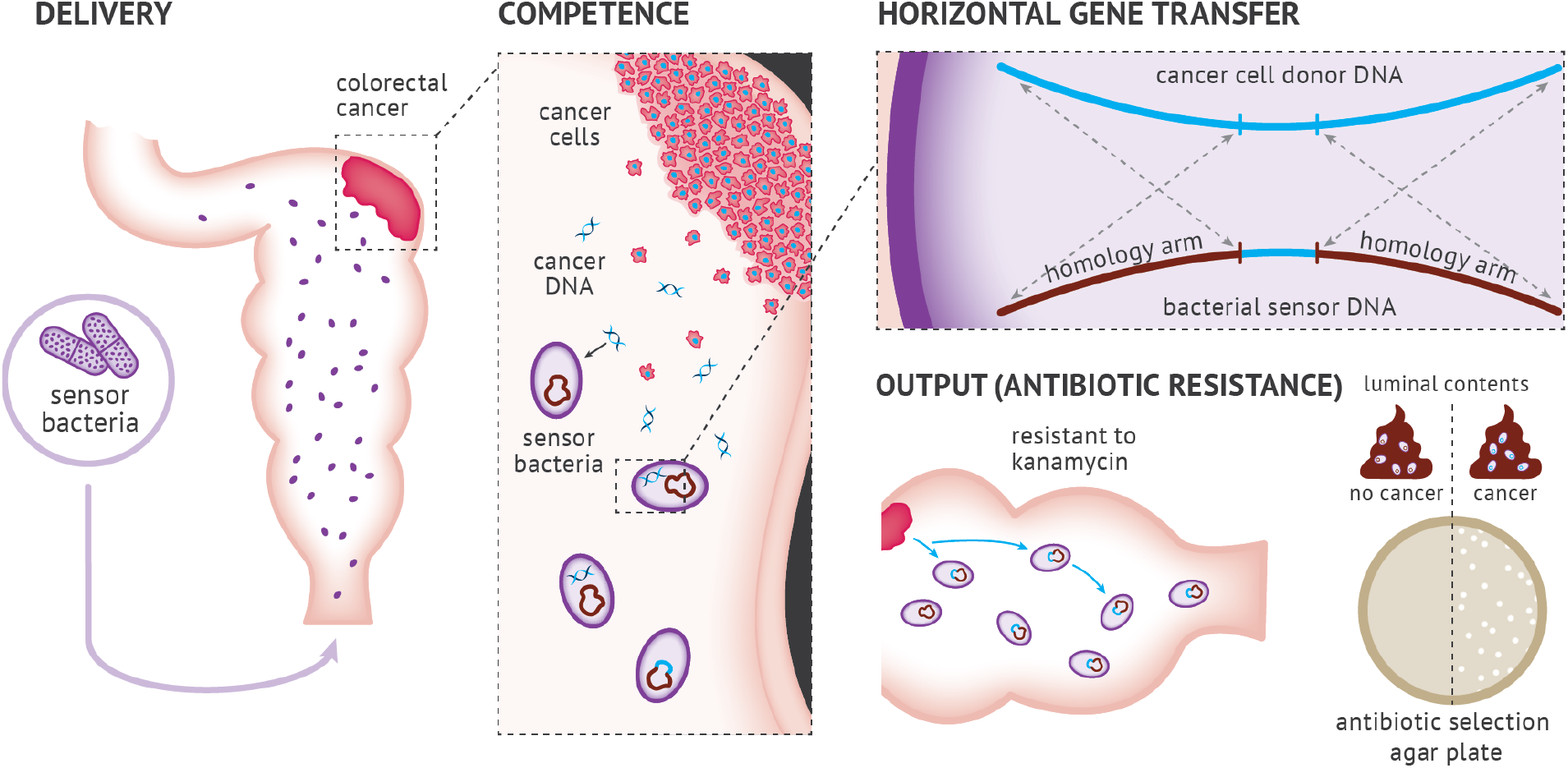
Engineered bacteria to detect tumor DNA. Engineered *A. baylyi* bacteria are delivered rectally in an orthotopic mouse model of CRC. The naturally competent *A. baylyi* take up tumor DNA shed into the colonic lumen. The tumor donor DNA is engineered with a *kan^R^* cassette flanked by *KRAS* homology arms (HA). The sensor bacteria are engineered with matching *KRAS* homology arms that promote homologous recombination. Sensor bacteria that undergo HGT from tumor DNA acquire kanamycin resistance and are quantified from luminal contents by serial dilution on antibiotic selection plates.

### Sensor bacteria can detect human cancer DNA

To test the hypothesis that bacteria could detect human tumor DNA, we generated transgenic donor human cancer cells and sensor bacteria (Fig. 2a). The donor cassette comprised a kanamycin resistance gene and GFP (*kan^R^-GFP*) flanked by 1 kb homology arms from human *KRAS* (Fig. 2b-c and Extended Data Fig. 1). *KRAS* is an important oncogene in human cancer, and a driver mutation in *KRAS* often accompanies the progression of simple into advanced colorectal adenomas^19^. We stably transduced this donor cassette into both RKO and LS174T human CRC cell lines using a lentiviral vector. To construct the sensor bacteria, we inserted a complementary landing pad with *KRAS* homology arms into a neutral genomic site of *A. baylyi*. We tested both a “large insert” design (2 kb), with a different resistance marker between the *KRAS* arms to be replaced by the donor cassette (Fig. 2b, Extended Data Fig. 2a), and a “small insert” design (8 bp), with the same *kan^R^-GFP* cassette as in the tumor donor DNA, but interrupted by 2 stop codons in *kan^R^* (Fig. 1 & 2c, Extended Data Fig. 2). The biosensor output was growth on kanamycin plates, measured as colony-forming units (CFUs) after serial dilution.

**Figure 2:**
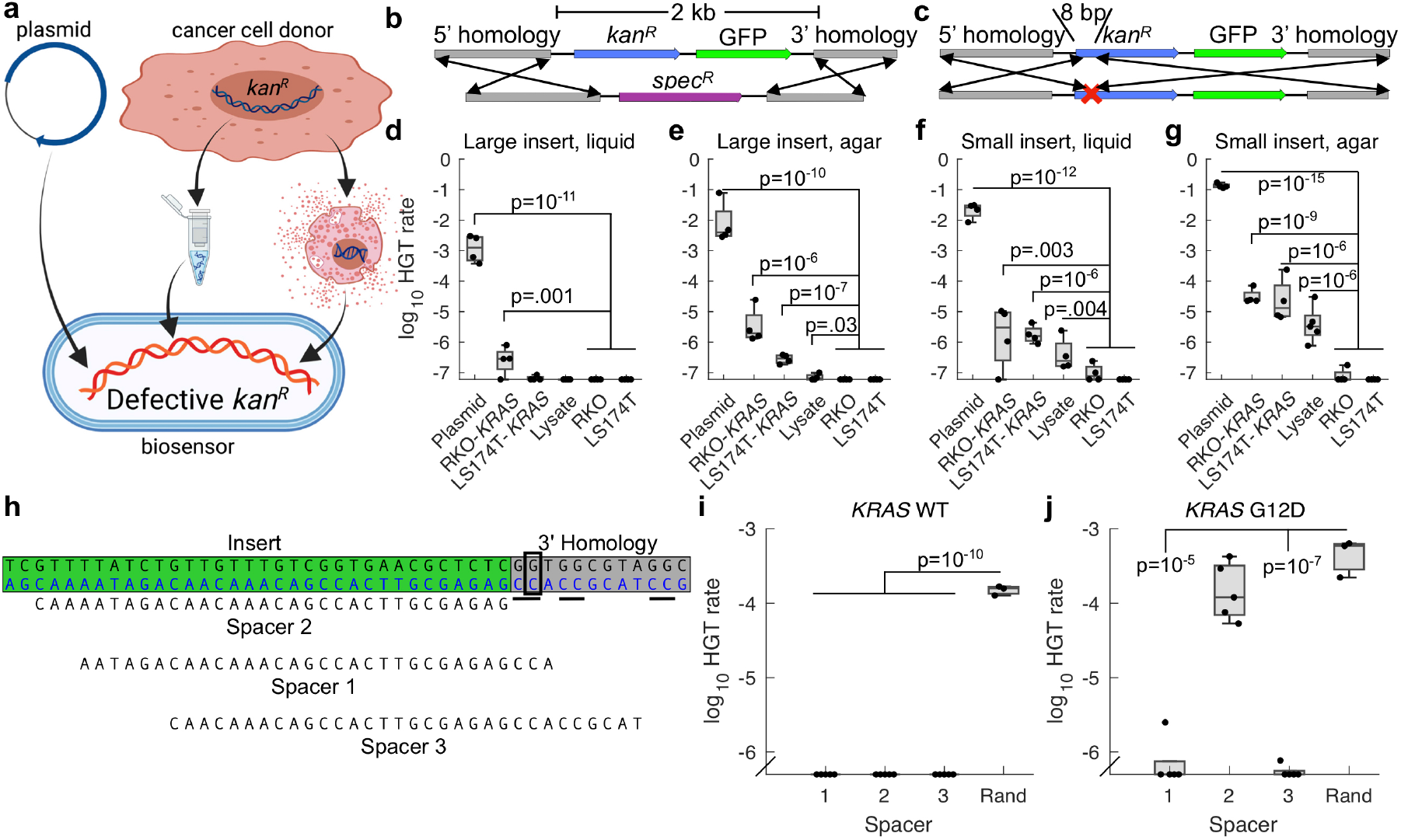
Sensing *KRASG12D* DNA *in vitro*. **a-c)** Donor DNA consisting of plasmid, purified cancer cell genomic DNA, or raw lysate (top) recombines into biosensor *A. baylyi* cells (bottom), transferring either a large, 2 kb insert (**b**), or a small, 8 bp insert to repair 2 stop codons (**c**), in both cases conferring kanamycin resistance. **d-g**) *A. baylyi* biosensors were incubated with plasmid DNA, purified RKO-*KRAS* or LS174T-*KRAS* genomic DNA, or raw RKO-*KRAS* lysate, all containing the donor cassette, or purified RKO or LS174T genomic DNA as controls. Biosensor cells included either “large insert” (**b,d,e**) or “small insert” (**c,f,g**) designs, and transformations were performed in liquid culture (**d,f)** or on solid agar surfaces (**e,g**). Two-sample t-tests compared data to combined RKO and LS174T genomic DNA controls for the same conditions. **h**) CRISPR spacers targeting the *KRAS* G12D mutation (boxed), using the underlined PAMs. **i,j** Fraction of total biosensor cells expressing the indicated CRISPR spacers that were transformed by plasmid donor DNA with wild type (**i**) or mutant G12D (**j**) *KRAS*. Statistics were obtained using two-sample, one-sided t-tests. Data points below detection are shown along the x-axis

We tested both designs using various donor DNA sources, both in liquid culture and on solid agar (Fig. 2a). The “large insert” biosensors detected donor DNA from purified plasmids and genomic DNA both in liquid (Fig. 2d) and on agar (Fig. 2e). On agar, they also detected raw, unpurified lysate, albeit at just above the limit of detection (Fig. 2e). As expected^20^, the “small insert” design improved detection efficiency roughly 10-fold, reliably detecting donor plasmid, purified genomic DNA, and raw lysate both in liquid and on agar (Fig. 2f-g, Extended Data Supplemental Movie). Across donor DNA and biosensor design, detection on solid agar was approximately 10-fold more efficient than in liquid culture. Importantly, detection of donor DNA from raw lysate demonstrated that the biosensors do not require *in vitro* DNA purification^21^.

*A. baylyi* can take up DNA at approximately 60 bp/s^22^. Given a human genome of 3.2 × 10^9^ bp, each *A. baylyi* cell, including its direct ancestors, can sample roughly 10^−3^ of a human genome in a 24-hour period. Combined with the data shown in Fig. 2g, with a detection rate around 10^−5^ per *A. baylyi* cell for RKO*-KRAS* and LS174T*-KRAS* donor DNA, this suggests a detection efficiency of around 1% per processed donor sequence. While this rough calculation assumes a constant DNA processing rate, the result is quite similar to what we found for HGT from *E. coli* to *A. baylyi*^21^.

### Sensor bacteria can discriminate wild-type from mutant *KRAS* DNA

Mutations in codon 12 of *KRAS* are present in 27% of CRC^23^, and are common in solid tumors generally^24^. To test whether sensor bacteria could discriminate between wild-type and mutant *KRAS (KRASG12D)*, which differ by a single G>A transition, we utilized *A. baylyi’s* endogenous Type I-F CRISPR-Cas system^25^. We stably transduced an RKO cell line with the *kan^R^-GFP* donor cassette flanked by wild-type *KRAS* (RKO-*KRAS*), and a second line with *KRASG12D* flanking sequences (RKO-*KRASG12D*). Next, we designed 3 CRISPR spacers targeting the wild-type *KRAS* sequence at the location of the *KRASG12D* mutation, using the *A. baylyi* protospacer-adjacent motif (PAM) of 5’-CC-protospacer-3’ (Fig. 2h). We inserted these as single-spacer arrays into a neutral locus in the “large insert” *A. baylyi* sensor genome.

The sensor bacteria, if effective, should reject wild-type *KRAS* through CRISPR-mediated DNA degradation. Conversely, the *KRASG12D* sequence should alter the target sequence and evade CRISPR-Cas interference. Two of the three spacers blocked transformation by both wild-type and mutant DNA (Fig. 2i-j). However, spacer 2, for which the *KRASG12D* mutation eliminated the PAM site, selectively permitted HGT only with *KRASG12D* donor DNA (Fig. 2i-j). The other common mutations in codon 12 of *KRAS* all eliminate this PAM as well^23^. Thus, sensor *A. baylyi* can be engineered to detect a mutational hotspot in the *KRAS* gene with single-base specificity.

### Sensor bacteria can integrate cancer DNA in organoid culture

*Ex vivo* organoid culture faithfully reflects endogenous tumor biology^26^. We therefore evaluated our sensor and donor constructs in organoid culture (Fig. 3a). We previously used CRISPR/Cas9 genome engineering to generate compound *Braf^V600E^; Tgfbr2^Δ/Δ^; Rnf43^Δ/Δ^; Znrf3^Δ/Δ^; p16Ink4a^Δ/Δ^* (BTRZI) mouse organoids that recapitulate serrated CRC when injected into the mouse colon^27^.

**Figure 3:**
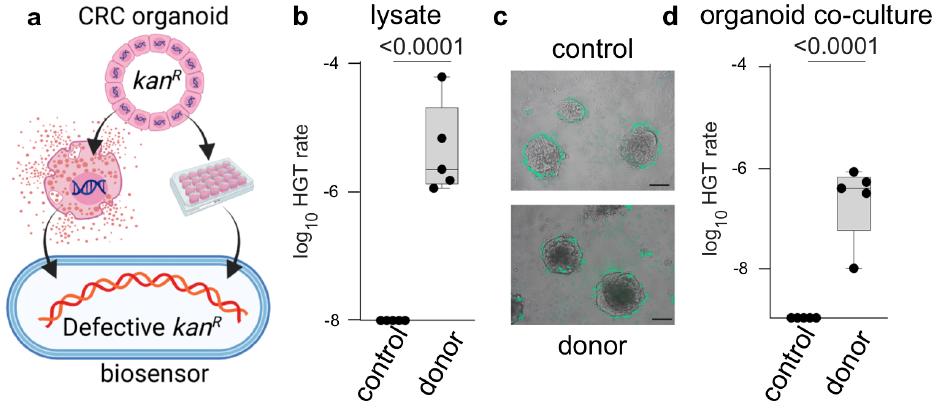
Detection of donor DNA from BTRZI-*KRAS-kan^R^* organoids. Schema depicting *in vitro* co-culture of *A. baylyi* sensor bacteria with BTRZI-*KRAS-kan^R^* (CRC donor) organoid lysates or viable organoids to assess HGT repair of kanamycin resistance gene (*kan^R^*). **b**. Recombination with DNA from crude lysates enables growth of *A. baylyi* sensor on kanamycin plates with transformation efficiency of 1.4×10^−5^ (limit of detection 10^−8^). **c**. Representative images of GFP-tagged *A. baylyi* sensor surrounding parental BTRZI (control) and BTRZI-*KRAS-kan^R^* donor organoids at 24h. Scale bar 100μm **d**. Co-culture of established CRC BTRZI-*KRAS-kan^R^* donor organoids with *A. baylyi* sensor enables growth of *A. baylyi* sensor on kanamycin plates with transformation efficiency 3.8×10^−7^ (limit of detection 10^−9^). In **b, d**, n = 5 independent experiments each with 5 technical replicates, one sample t-test on transformed data was used for statistical analysis with *P* values as indicated.

We transduced BTRZI organoids with the human *KRAS-* flanked donor DNA construct (*KRAS-kan^R^*) to generate donor CRC organoids and incubated their lysate with the more efficient “small insert” *A. baylyi* biosensors. As with the CRC cell lines, the sensor *A. baylyi* incorporated DNA from donor organoid lysate, but not from control lysates from the parental organoids (Fig. 3b, Extended Data Fig. 3a). Next, we co-cultured GFP-expressing sensor *A. baylyi* with BTRZI parental or BTRZI-*KRAS-kan^R^* donor organoids for 24 hours on Matrigel. The GFP-expressing sensor bacteria surrounded the organoids (Fig. 3c). Following co-culture with donor, but not parental, organoids, the *A. baylyi* sensor bacteria acquired donor DNA via HGT (Fig. 3d). HGT of kanamycin resistance was confirmed by Sanger sequencing of individual colonies (Extended Data Fig. 3c). Note that these experiments did not test specificity for mutant *KRAS*, but whether organoid-to-bacteria HGT would occur in organoid co-culture.

### Sensor bacteria can detect tumor DNA *in vivo*

Given that cancer-to-bacterial HGT occurred *in vitro*, both in cell lines and in organoid co-culture, we sought to test the CATCH system *in vivo*. We first confirmed that our BTRZI, orthotopic CRC model released tumoral DNA into the fecal stream. In this mouse model of CRC, engineered CRC organoids were injected orthotopically, by mouse colonoscopy, into the mouse colon to form colonic tumors, as previously described^27^. Using digital droplet PCR, we measured *Braf* mutant tumor DNA in stools collected from tumor-bearing and control mice. The BTRZI model reliably released tumor DNA into the colonic lumen (Extended Data Fig. 4).

We next conducted an orthotopic CRC experiment (Fig. 4a). NSG mice were injected colonoscopically with donor BTRZI-*KRAS-kan^R^* or non-donor BTRZI organoids or neither. All study groups were housed in separate cages. At week 5, once the tumors had grown into the lumen, 4×10^10^ sensor (or parental) *A. baylyi* bacteria were delivered via rectal enema. 24 hours later, a second enema of sensor or control bacteria were again administered. The mice were subsequently sacrificed and the colorectum harvested with the luminal effluent plated for analysis. Sensor bacteria contained an additional chloramphenicol resistance gene to aid detection, and previous experiments had confirmed that a combination of vancomycin, chloramphenicol and kanamycin provided the best selection for discriminating biosensor HGT from other resistant fecal microbiota. Serial dilutions were plated on agar with different antibiotic combinations for the quantification of HGT in both biosensor and parental *A. baylyi* (Fig 4b).

**Figure 4.**
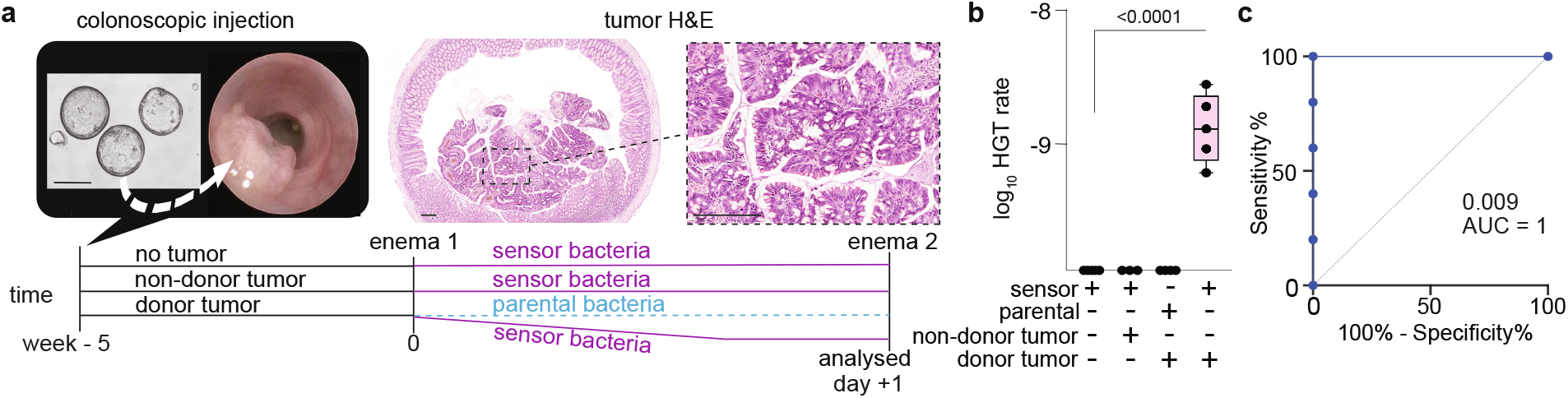
Horizontal gene transfer detected in luminal contents from mice bearing BTRZI-KRAS-kanR tumors after rectal dosing of *A. baylyi* sensor bacteria. **a**, Schema depicting *in vivo* HGT experiments: generation of BTRZI-KRAS-kanR (CRC donor) tumors in mice via colonoscopic injection of CRC donor organoids with tumor pathology validated by H&E histology, administration of sensor *A. baylyi* or parental *A. baylyi* and analysis of luminal contents. Scale bars 200μm. **b**, rectal delivery of *A. baylyi* sensor to mice bearing CRC donor tumors results in kanamycin resistant *A. baylyi* sensor in luminal contents via HGT with transformation efficiency of 4.75×10^−9^ (limit of detection 1.25×10^−10^). HGT rate calculated from CFU on Kanamycin/Chloramphenicol/Vancomycin (transformants) and Chloramphenicol/Vancomycin (total A. baylyi) selection plates, n=3-5 mice/group. One-way Anova with Tukey’s post-hoc on log_10_ transformed data was used for statistical analysis with P values shown. **c**, ROC curve analysis of HGT CFU following enema, AUC = 1, p = 0.009.

Following sensor bacteria delivery, the kanamycin-resistant CFUs were only present in the donor tumor-bearing mice that were administered sensor bacteria. There was no HGT detected in any control groups (Fig. 4b). The resistant colonies were confirmed to be the engineered biosensor strain by antibiotic resistance, green fluorescence, 16S sequencing, and HGT-mediated *kan^R^* repair of individual colonies (Extended Data Fig. 5). Our CATCH system perfectly discriminated mice with and without CRC (Fig 4c). Unfortunately, *A. baylyi* biosensors did not establish a sufficient colonising population within the colon to detect tumor bearing mice from collected stool (Extended data Fig. 6). The biosensor population only reached 10^5^ per stool which, based on our *in vitro* studies, is insufficient to detect HGT with the current system (Fig 3d).

### Sensor bacteria can detect and genotype natural, non-engineered, tumor DNA

Finally, we designed living biosensors that can detect and analyze non-engineered DNA without a donor cassette. The *tetR* repressor gene was inserted between the *KRAS* homology arms in the biosensor, and in a second locus, we placed an output gene under control of the P_LtetO-1 promoter^28^ (Fig. 5a). Here, the output gene was kanamycin resistance for ease of measurement, but the output gene is arbitrary and readily changed. Interestingly, we found that wild type *tetR* is toxic in *A. baylyi*, but we fortuitously isolated a temperature sensitive mutant^29^ that did not kill the biosensors even at the permissive temperature (see Methods). Clones containing wild type *tetR* could only be isolated in the presence of the inducer anhydrotetracycline (aTc), which inactivates *tetR*, and they could not grow without aTc. We hypothesize that wild type *tetR* disrupts essential gene expression by binding to off-target sites in the *A. baylyi* genome, and that the temperature sensitive mutation also destabilizes binding to off-target sequences, thus increasing binding specificity.

**Figure 5:**
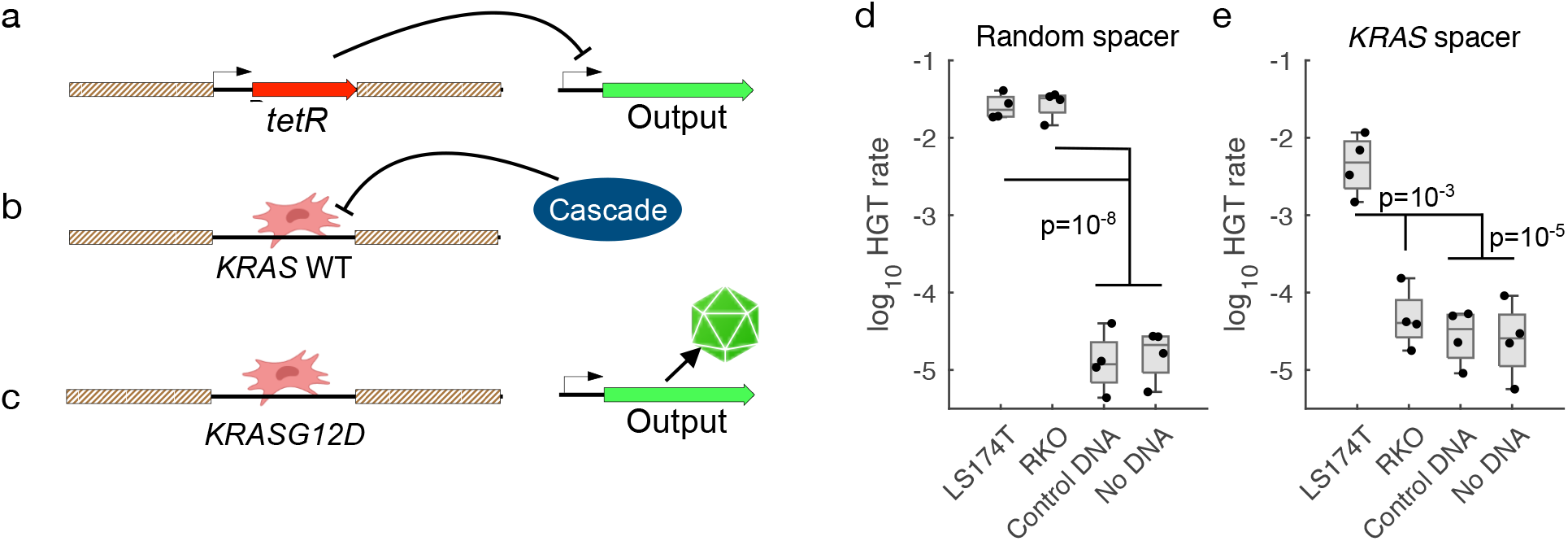
Detection of non-engineered DNA. **a**, Prior to recombination with target DNA, *tetR* is located between the homology arms on the *A. baylyi* genome and expression of the output gene is repressed. **b**, Target DNA with the *KRAS* homology arms and wild-type sequence is recognized and degraded by the Type I-F CRISPR-Cas effector complex, Cascade. **c**, Target DNA with the *KRASG12D* mutation avoids degradation by Cascade, replaces *tetR* in the biosensor genome, and relieves repression of the output gene. **d-e**, Fraction of biosensors with either a random CRISPR spacer (**d**) or a spacer targeting wild type *KRAS* (**e**) that detected donor DNA sequences PCRed from LS174T or RKO cells, unrelated plasmid DNA, or no DNA. Statistics were obtained using 2-sample t-tests.

In this design, expression of the output gene is constitutively repressed (Fig. 5a). Upon recombination with the target DNA, the repressor *tetR* is deleted from the genome and replaced with *KRAS* donor DNA from human cells. If the *KRAS* sequence is wild type at the G12 locus, Cascade, the Type I-F CRISPR-Cas effector complex, detects and degrades it (Fig. 5b). However, if the G12 locus is mutated, the PAM site and therefore CRISPR-Cas targeting are eliminated, and expression from the output gene turns on (Fig. 5c).

We tested this natural DNA sensor design *in vitro* using PCR products from LS174T and RKO cells as donor DNA. Natural DNA biosensors with a random CRISPR spacer detected DNA sequences from both cell lines (Fig. 5d), and biosensors with the *KRAS* spacer accurately detected only DNA sequence from LS174T cells, which contain the *KRASG12D* mutation, demonstrating biosensor detection and discrimination of natural target DNA.

## Discussion

In this study, naturally competent *A. baylyi* were engineered to sense donor DNA from cancer cells. Our CATCH biosensor system was optimized *in vitro* and then validated *in vivo* using an orthotopic mouse model of CRC. Furthermore, we engineered a CRISPR-based technique to provide specificity for the mutant *KRASG12D* vs. wild-type *KRAS* for both engineered and natural tumor sequences. The sensor bacteria described here demonstrate that a living biosensor can detect tumor DNA shed from CRC *in vivo* in the gut, with no sample preparation or processing. The sensor is highly sensitive and specific, with 100% discrimination between mice with and without CRC. Importantly, engineered donor cassettes are not required for biosensors to detect, discriminate, and report on target sequences, although these natural DNA sensors will need an improved signal-to-background ratio to reliably detect sequences within whole genomic DNA. The homology arms and CRISPR spacers are modular, so this strategy could readily be generalized to detect and analyze arbitrary target sequences of interest.

*In vitro* DNA analysis helps detect and manage important human diseases, including cancer and infection^30^. However, *in vitro* sensing requires potentially invasive removal of samples, and many DNA diagnostics cannot achieve clinically relevant sequence resolution, with more advanced techniques remaining too expensive for routine use in all settings^31^. Direct sampling of the gut *in vivo* may offer important advantages. The gastrointestinal tract contains significant DNase activity^32^, which limits the lifetime of free DNA in both rodents and humans^18,33,34^, and may thus reduce the information content of downstream fecal samples^35–37^. Bacterial biosensors located *in situ* could capture and preserve DNA shortly after its release, before degradation by local DNases. Perhaps the most exciting aspect of CATCH, however, is that unlike *in vitro* diagnostics, once target DNA is captured, it could be coupled to direct and genotype-complementary delivery of nanobodies, peptides, or other small molecules for the treatment of cancer or infection^38,39^.

To realise these opportunities and to translate this technology into the detection and management of human disease, however, the CATCH system will need further development. For *in vivo* applications, biosensors other than *A. baylyi* may be needed that are more compatible with the specific diagnostic niche of interest. Safety and biocontainment would also need to be ensured before use in humans, which could be addressed through *e.g*. genomic recoding to introduce synthetic auxotrophy, which could prevent unwanted HGT and growth^40,41^. Furthermore, dosing and conditioning regimens are needed to optimise engraftment of biosensors. However, mice are far smaller than humans, and a larger scale would likely benefit *in vivo* CATCH by increasing both the number of biosensors and the released target DNA. Finally, new bioengineering to amplify target DNA through HGT-mediated selection, intercellular quorum sensing circuits, or intracellular genetic memory switches would all help to apply this work^9,11^.

CATCH is a new approach for detecting, diagnosing, and in the future potentially treating disease. Furthermore, CATCH may be useful in many non-clinical applications as well, wherever genetic detection is important, continuous surveillance is desirable, or an immediate and localized biologically-generated response would be beneficial.

## Supporting information

Supplemental DNA

## Methods

### Data availability

All data generated or analyzed during this study are included in this published article (and its supplementary information files), and raw data files are available upon request.

### Bacterial cell culture and cloning to generate biosensors

*Acinetobacter baylyi* ADP1 was obtained from the American Type Culture Collection (ATCC #33305) and propagated in standard LB media at 30 or 37 °C. *KRAS* homology arms were inserted into a neutral genetic locus denoted *Ntrl1*, replacing the gene remnant ACIAD2826. For the “large insert” design, a spectinomycin resistance gene was placed between the *KRAS* homology arms. For the “small insert” design, two stop codons were placed near the beginning of the *kan^R^* gene of the donor cassette, and the broken cassette was inserted into *A. baylyi*. CRISPR arrays were inserted into a neutral locus used previously, replacing ACIAD2186, 2187 and part of 2185. Ectopic CRISPR arrays were driven by a promoter region that included 684 bp from upstream of the first repeat of the endogenous, 90-spacer array.

For natural DNA biosensors, a temperature-sensitive *tetR* repressor was placed between the *KRAS* homology arms. An output gene, either *kan^R^* or GFP, was placed under control of the P_LtetO-1 promoter in a second neutral locus denoted *Ntrl2*, replacing the gene remnants ACIAD1076-1077. Repeated attempts to clone wild type *tetR* into *A. baylyi* failed, but we fortuitously isolated a temperature sensitive mutant with two mutations: W75R and an additional 8 amino acids on the C terminus. This mutant *tetR* permitted growth at both 30 and 37 °C, but it only repressed its target at 30 °C. The W75R mutant had been isolated previously in an intentional screen. We were able to clone wild-type *tetR* on the inducer aTc, but it was unable grow without aTc at any temperature.

### *In vitro* biosensor transformation experiments

*A. baylyi* were grown overnight in LB at 30 °C. Cells were then washed, resuspended in an equal volume of fresh LB, and mixed with donor DNA. For transformation in liquid, 50 μl cells were mixed with 250 ng donor DNA and incubated in a shaker at 30 °C for 2 hours or overnight. For transformation on agar, 2 μl cells were mixed with >50 ng donor DNA, spotted onto LB plates containing 2% wt/vol agar, and incubated at 30 °C overnight. Spots were cut out the next day and resuspended in 500 μl phosphate buffered saline solution (PBS). To count transformants, cells were 10-fold serially diluted 5 times, and 2 μl spots were deposited onto selective (30 ng/ml kanamycin) and non-selective 2% agar plates, with 3 measurement replicates at each dilution level. Larger volumes of undiluted samples were also spread onto agar plates to increase detection sensitivity (25μl for liquid culture, 100 μl for resuspended agar spots). Colonies were counted at the lowest countable dilution level after overnight growth at 30 °C, and measurement replicates were averaged. Raw, unpurified lysate was produced by growing donor RKO cells in a culture dish until confluence, trypsinizing and harvesting cells, pelleting them in a 15 ml tube, resuspending them in 50 μl PBS, and placing the tube in a –20 °C freezer overnight to disrupt cell membranes.

### *In vitro* **statistics**

Hypothesis testing was performed using 2-sample, one-sided t-tests in Matlab after taking base 10 logarithms, since serial dilutions produce log-scale data. Where data points were below the limit of detection, they were replaced by the limit of detection as the most conservative way to include them in log-scale analysis. Comparisons between large vs small inserts or liquid vs solid agar culture were performed using paired t-tests, where data were matched for donor DNA and either culture type (liquid vs agar) or insert size, respectively. For Figure 2, d-g) n=4, i,j) n=5 except for random spacer n=3.

### Creation of RKO and LS174T donor cell lines

To create RKO donor and LS174T donor cell lines, lentiviral expression plasmid pD2119- FLuc2 KRasG12D donor was co-transfected with viral packaging vectors, psPAX2 (Addgene; plasmid; 12260) and MD2G (Addgene; plasmid; 12259), into HEK293T cells. At 48 and 72 h after transfection, viral supernatants were harvested, filtered through a 0.45-μm filter, and concentrated using Amicon Ultra Centrifugal Filters (Merck Millipore; UFC910024). Concentrated lentivirus particles were used for transduction. The viral supernatant generated was used to transduce RKO and LS174T cells. 48 hours after transduction, stable transformants were selected with 4 μg/ml puromycin. Cell lines identity was confirmed by STR analysis. KRAS status of RKO (KRAS wildtype) and LS174T (KRAS G12D) cell lines was confirmed by amplification of a 220bp PCR fragment of the exon 2 KRAS gene, including codons 12 and 13 with primers KRAS F: GGTGGAGTATTTGATAGTGTATTAACC and KRAS R: AGAATGGTCCTGCACCAGTAA. Sanger sequencing was conducted using the same primers.

### Creation of BTRZI CRC donor organoids

BTRZI (Braf^V600E^;Tgfbr2^Δ/Δ^;Rnf43 ^Δ/Δ^ /Znf43 ^Δ/Δ^;p16 Ink4a ^Δ/Δ^) organoids were generated using CRISPR-Cas9 engineering^27^ and grown in 50 μl domes of GFR-Matrigel (Corning,; 356231) in organoid media: Advanced Dulbecco’s modified Eagle medium/F12 (Gibco; 12634010) supplemented with 1x gentamicin/antimycotic/antibiotic (Gibco; 15710064, 15240062), 10mM HEPES (Gibco; 15630080), 2 mM GlutaMAX (Gibco; 35050061), 1x B27 (Gibco; 12504-044), 1x N2 (Gibco; 17502048), 50 ng/ml mouse recombinant EGF (Peprotech; 315-09), 10 ng/ml human recombinant TGF-β1 (Peprotech; 100-21). Following each split, organoids were cultured in 10 μM Y-27632 (MedChemExpress; HY-10583), 3 μM iPSC (Calbiochem; 420220), 3 μM GSK-3 inhibitor (XVI, Calbiochem; 361559) for the first 3 days.

To create BTRZI CRC donor organoids, lentiviral expression plasmid pD2119-FLuc2 KRasG12D donor was co-transfected with viral packaging vectors, psPAX2 (Addgene; plasmid; 12260) and MD2G (Addgene; plasmid; 12259), into HEK293T cells. At 48 and 72 h after transfection, viral supernatants were harvested, filtered through a 0.45-!m filter, and concentrated using Amicon Ultra Centrifugal Filters (Merck Millipore; UFC910024). Concentrated lentivirus particles were used for transduction. The viral supernatant generated was used to transduce BTRZI organoids by spinoculation. Briefly, organoids were dissociated to single cells using TrypLE. 1×10^5^ single cells were mixed with 250 μl organoid media; 10 μM Y-27632; 250 μl concentrated viral supernatant and 4 μg/ml polybrene (Sigma,; H9268) in a 48 well tray before centrifugation at 600 xg for 90 minutes at 32 °C. Meanwhile, 120 μl 50:50 ADMEM:Matrigel mixture was added to a cold 24-well tray before centrifugation of this bottom matrigel layer for 40 minutes at 200xg at room temperature, followed by solidifying the Matrigel by incubating at 37 °C for 30 minutes. After spinoculation, cells were scraped from the well and plated on top of the Matrigel monolayer with organoid media. The following day, the media was removed and the upper layer of Matrigel was set over the organoids by adding 120 μl 50:50 ADMEM:Matrigel and allowing to set for 30 minutes before adding organoid media. 48 hours after transduction, BTRZI donor organoids were selected with 8 μg/ml puromycin for 1 week, then maintained in organoid media with 4 μg/ml puromycin.

### Organoid lysate mixed with *A. baylyi* sensor bacteria

BTRZI (parental) and BTRZI donor organoids were grown for 5 days in 50 ml Matrigel domes. Organoids were dissociated to single cells with TrypLE, counted and 6×10^5^ single cells were collected in PBS and snap frozen. The CFU equivalence of exponentially growing *A. baylyi* sensor culture at OD_600_ 0.35 was ascertained by serial dilution of 3 independent cultures with 5 technical replicates plated on 10 μg/ml Chloramphenicol LB agar plate to be 2.4 × 10^8^ CFU per ml. *A. baylyi* sensor was grown in liquid culture with 10 μg/ml Chloramphenicol to OD_600_ 0.35 before mixing with organoid lysate at a 1:1 ratio and grow overnight on LB agar plates at 30 °C. All bacteria was scraped into 200 μl LB/20% glycerol before spotting 5× 5 μl spots onto kanamycin and chloramphenicol plates and grown overnight at 37 °C. Colonies were counted and the dilution factor was accounted for to calculate CFU per ml. Rate of HGT was calculated by dividing the CFU per ml of transformants (Kanamycin plates) by the CFU per of total *A. baylyi* (chloramphenicol plates) for 5 independent experiments.

### Coculture organoids with *A. baylyi* sensor bacteria

For co-culture experiments, 24-well trays were coated with Matrigel monolayers. Briefly, 200 μl 50:50 ADMEM:Matrigel mixture was added to a cold 24-well tray and centrifuged for 40 minutes at 200xg at room temperature, followed by a 30 minute incubation at 37 °C to solidify matrigel. BTRZI (parental) and BTRZI donor organoids were dissociated into small clusters using TrypLE and grown for 5 days on a Matrigel monolayer in organoid media without antibiotics before 50 μl OD_600_ 0.35 *A. baylyi* sensor was added to each well. After 24 hours, organoids were photographed then collected and grown overnight on LB agar plates at 30 °C. All bacteria was scraped into 200 μl LB/20% glycerol before spotting 5x 5 μl spots onto kanamycin and chloramphenicol plates and grown overnight at 37 °C. Colonies were counted and the dilution factor was accounted for to calculate CFU per ml. Rate of HGT was calculated by dividing the CFU per ml of transformants (kanamycin plates) by the CFU per ml of total *A. baylyi* (chloramphenicol plates) for 5 independent experiments.

### Horizontal gene transfer *in vivo*

BTRZI donor organoids were isolated from Matrigel and dissociated into small clusters using TrypLE. The cell clusters (equivalent to ~150 organoids per injection) were washed three times with cold PBS containing 10 μM Y-27632 and then resuspended in 20 μl 10% GFR matrigel 1:1000 indian ink, 10 μM Y-27632 in PBS and orthotopically injected into the mucosa of the proximal and distal colon of anaesthetised 10-13 week old NSG mice (150 organoids per injection), as previously described^27^. Briefly, a customised needle (Hamilton Inc. part number 7803-05, removable needle, 33 gauge, 12 inches long, point 4, 12 degree bevel) was used. In each mouse up to 2 injections of 20μl were performed. CRC donor tumor growth was monitored by colonoscopy for 5 weeks and the videos were viewed offline using QuickTime Player for analysis. Colonoscopy was performed using a Karl Storz Image 1 Camera System comprised of: Image1 HDTV HUB CCU; Cold Light Fountain LED Nova 150 light source; Full HD Image1 3 Chip H3-Z Camera Head; Hopkins Telescope, 1.9mm, 0 degrees. A sealed luer lock was placed on the working channel of the telescope sheath to ensure minimal air leakage (Coherent Scientific, # 14034-40). Tumor growth of the largest tumor visualised was scored as previously described using the Becker Scale (Rex et al, 2012 Am J Gastroenterol). *A. baylyi* sensor was grown in LB liquid culture with 6 μg/ml Chloramphenicol to OD_600_ 0.6. *A.baylyi* parental was grown in LB liquid culture to OD_600_ 0.6. *A. baylyi* was washed twice with PBS before 13 mice received 4×10^10^ *A. baylyi* sensor via enema (5 mice without tumors; 3 mice with non-donor BTRZI CRC tumors and 5 mice with BTRZI CRC donor tumors), 4 mice received 4×10^10^ *A. baylyi* parental via enema. Enema was performed as per previous publication. Briefly, mice were anaesthetised with isofluorane and colon flushed with 1 ml of room temperature sterile PBS to clear the colon cavity of any remaining stool. A P200 pipette tip coated with warm water was then inserted parallel into the lumen to deliver 50 μl of bacteria into the colon over the course of 30 seconds. After infusion, the anal verge was sealed with Vetbond Tissue Adhesive (3M; 1469SB) to prevent luminal contents from being immediately excreted. Animals were maintained on anaesthesia for 5 minutes, and then allowed to recover on heat mat and anal canal inspected 6 hours after the procedure to make sure that the adhesive has been degraded. 24 hours after *A. baylyi* administration, mice received a second enema dosing. Mice were then culled, colons were removed and luminal contents were collected. Luminal contents were grown overnight at 37 °C on LB agar with 10 μg/ml vancomycin plates. All bacteria was collected into 250 μl LB/20% glycerol, vortexed and stored at −80 °C. 5x 5μl serial dilutions were spotted onto LB agar plates containing (1) vancomycin (to detect total *A.baylyi* parental); (2) chloramphenicol; vancomycin (to detect total *A.baylyi* sensor) and (3) kanamycin; chloramphenicol; vancomycin (to detect recombined *A.baylyi* sensor). Colonies were counted and dilutions were factored to calculate CFU *A. baylyi* per mouse. For experiments analysing *A.baylyi* in stool, BTRZI CRC donor tumors were established and monitored as described above. After 5 weeks of tumor growth, 9 mice received *A.baylyi* sensor enemas (5 mice without tumors; 4 mice with BTRZI CRC donor tumors) and 6 mice received *A.baylyi* parental enemas (3 mice without tumors and 3 mice with BTRZI CRC donor tumors). Stool was collected 24 hours after *A. baylyi* administration into 250 μl PBS/20% glycerol, vortexed and stored at −80 °C. Stool was analysed on LB agar plates containing (1) vancomycin (to detect total *A.baylyi* parental); (2) chloramphenicol; vancomycin (to detect total *A.baylyi* sensor) and (3) kanamycin; chloramphenicol; vancomycin (to detect recombined *A.baylyi* sensor). Colonies were counted and dilutions were factored to calculate CFU *A. baylyi* per mouse.

### Sequencing gDNA from bacterial colonies grown on kanamycin plates

*A. baylyi* transformants were individually picked from kanamycin; vancomycin plates and grown in liquid culture LB supplemented with 25 μg/ml kanamycin, 10 μg/ml vancomycin and 6 μg/ml chloramphenicol. gDNA was extracted using purelink genomic DNA minikit (Invitrogen; K182001). Genomic regions of interest were amplified using Primestar Max DNA polymerase (Takara, # R045A) and primers HGTpcrF: CAAAATCGGCTCCGTCGATACTA; HGTpcrR: TAGCATCACCTTCACCCTC and 16S 27Fa: AGAGTTTGATCATGGCTCAG; 16s 27Fc: AGAGTTTGATCCTGGCTCAG; 16S 1492R: CGGTTACCTTGTTACGACTT (16S 27Fa:16S 27Fc: 16S 1492R = 0.5:0.5:1).Sanger sequencing was conducted using the same primers.

## Acknowledgements

Mr Phil Winning for design assistance with Figure 1. This work was supported by NIH grant R01CA241728. Professor Barbara Leggett and A/Prof. Vicki Whitehall for the original gift of the parental RKO and LS174T human CRC cell lines used in this study.

## Author contributions

RC, DW & JH conceived of the concept and study plan. RC, JW, JN, JG, NS, YL, MI, GR, FR, SK, ET, LV, SW, DW, & JH were all involved with data acquisition and or interpretation. RC, JW, RK, SW, DW, & JH were involved in writing and revising the final manuscript.

## Competing interest declaration

J.H. is a co-founder and board member with equity in GenCirq Inc, which focuses on cancer therapeutics.

Reprints and permissions information is available at www.nature.com/reprints

## Extended Data

**Extended Data Figure 1**: Plasmid donor DNA used to transfect mammalian cell lines and as positive control donor DNA for *in vitro* experiments.

**Extended Data Figure 2**: “Large insert” (**a**) and “small insert (**b**) designs for the biosensors. *KRAS* homology arms are shown in striped gray with surrounding genomic context outside them. Note that large and small inserts refers to the size of the donor DNA region that must transfer to confer kanamycin resistance, not to the size of the region between homology arms in the biosensor. Two single-base changes introducing nearby stop codons at the beginning of *kan^R^* are shown for the small insert design (**b**).

**Extended Data Figure 3: Sensor detection of donor DNA from BTRZI CRC organoids** *A. baylyi* sensor bacteria are constitutively chloramphenicol resistant, hence c*hlorR* CFUs provide a read-out of total *A. baylyi* present. In contrast, kanamycin resistant sensor bacteria rely on incorporation of donor DNA from CRC organoids to correct the defective *kan* gene and enable growth on kanamycin selection plates. **a** Recombination with lysate from CRC donor organoids enables growth of *A. baylyi* sensor on kanamycin plates. Shown here with representative plates and CFU analysis. **b** After co-culturing established CRC donor organoids with *A. baylyi* sensor, recombination with donor DNA from CRC donor organoids enables growth of *A. baylyi* sensor on kanamycin plates. Shown here with representative images and CFU analysis. Scale bars 200 μm. **a, b**, Fig 3 contains the same data as shown here but presented as HGT rate (kanamycin resistant CFU *A.baylyi* per ml/chloramphenicol CFU *A.baylyi* per ml), n = 5 independent experiments each with 5 technical replicates. **c** Representative Sanger sequencing chromatograms of PCR amplicon covering the region of the *kan* gene containing informative SNPs, to highlight the difference in sequence in gDNA isolated from parental *A. baylyi* sensor bacteria compared to *A.baylyi* colonies isolated from kanamycin plates following mixing with donor organoid lysates or viable organoids.

**Extended Data Figure 4: High sensitivity digital droplet PCR (ddPCR) detection of CRC mutation (***BrafV600E***) in stool DNA isolated from tumour bearing animals (n=3-4 mice/group). a**, Representative images of ddPCR data. **b**, CRC mutation (*BrafV600E)* positive droplets as a % of total droplets. Analysis of no template negative control samples and stool DNA samples from non-tumour bearing animals was used to determine the sensitivity threshold of the assay. Positive control samples contain 10% *BrafV600E* gDNA spiked into stool DNA sample from non-tumour bearing animal. NT, no tumour; Ts, small tumour; Tm, medium tumour; Tl, large tumour; NTC, no template PCR negative control.

**Extended Data Figure 5**:Horizontal gene transfer is detected in luminal contents from mice bearing BTRZI CRC donor tumors after rectal dosing of *A. baylyi* sensor bacteria. a Recombined *A. baylyi* transformants are GFP positive on kanamycin/chloramphenicol/vancomycin selection plates, scale bar 500 !m. b Representative Sanger sequencing chromatograms of PCR amplicon covering the region of the kan gene containing informative SNPs to highlight the difference in sequencing DNA isolated from parental *A. baylyi* sensor bacteria compared to *A. baylyi* colonies isolated from kanamycin/chloramphenicol/vancomycin plates in luminal contents from mice bearing BTRZI CRC donor tumors after rectal dosing of *A. baylyi* sensor bacteria

**Extended data figure 6**:Horizontal gene transfer is not detected in stool from mice bearing BTRZI CRC donor tumors after rectal dosing of *A. baylyi* sensor bacteria. **a**, Schema depicting *in vivo* HGT experiment: generation of BTRZI-KRAS-kanR (CRC donor) tumors in mice via colonoscopic injection of CRC donor organoids with tumor pathology validated by H&E histology, administration of parental or sensor *A. baylyi* and stool collection. **b,c** Rectal delivery of *A. baylyi* sensor to mice bearing CRC tumors results in no detection of HGT in *A. baylyi* sensor bacteria present within stool. Data points represent the average CFU per stool from 4 stools per mouse grown on b kanamycin/vancomycin selection plates (*A. baylyi* sensor HGT) or c chloramphenicol/vancomycin selection plates (total A.baylyi sensor), n=3-5 mice/group. Limit of detection 80 CFUs.

**Extended Data Movie 1**: *A. baylyi* biosensors taking up plasmid donor DNA.

*A. baylyi* were grown overnight, washed into fresh LB, mixed with saturating pLenti-KRAS donor DNA, and sandwiched between an agar pad and a glass bottom dish. Images were taken every 10 minutes. GFP fluorescence indicates that the cells have taken up and genomically integrated the donor DNA cassette.

**Extended Data DNA Files**:

DNA cassettes and surrounding regions corresponding to the “large insert”, “small insert”, and natural DNA sensor designs for *A. baylyi*, and the plasmid donor DNA, as shown in Extended Data 1,2, in Genbank format.

