## Extended Data Figure for "Engineered bacteria detect tumor DNA"

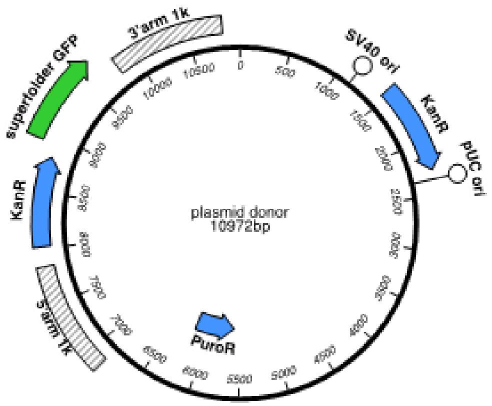


**Extended Data Figure 1.** Plasmid donor DNA used to transfect mammalian cell lines and as positive control donor DNA for *in vitro* experiments.

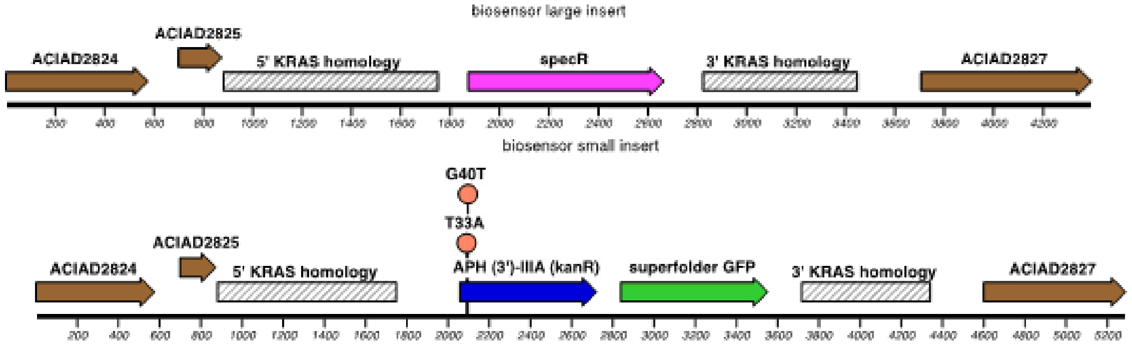


**Extended Data Figure 2.** "Large insert" (**a**) and "small insert (**b**) designs for the biosensors. *KRAS* homology arms are shown in striped gray with surrounding genomic context outside them. Note that large and small inserts refers to the size of the donor DNA region that must transfer to confer kanamycin resistance, not to the size of the region between homology arms in the biosensor. Two single-base changes introducing nearby stop codons at the beginning of *kan*$^R$ are shown for the small insert design (**b**).

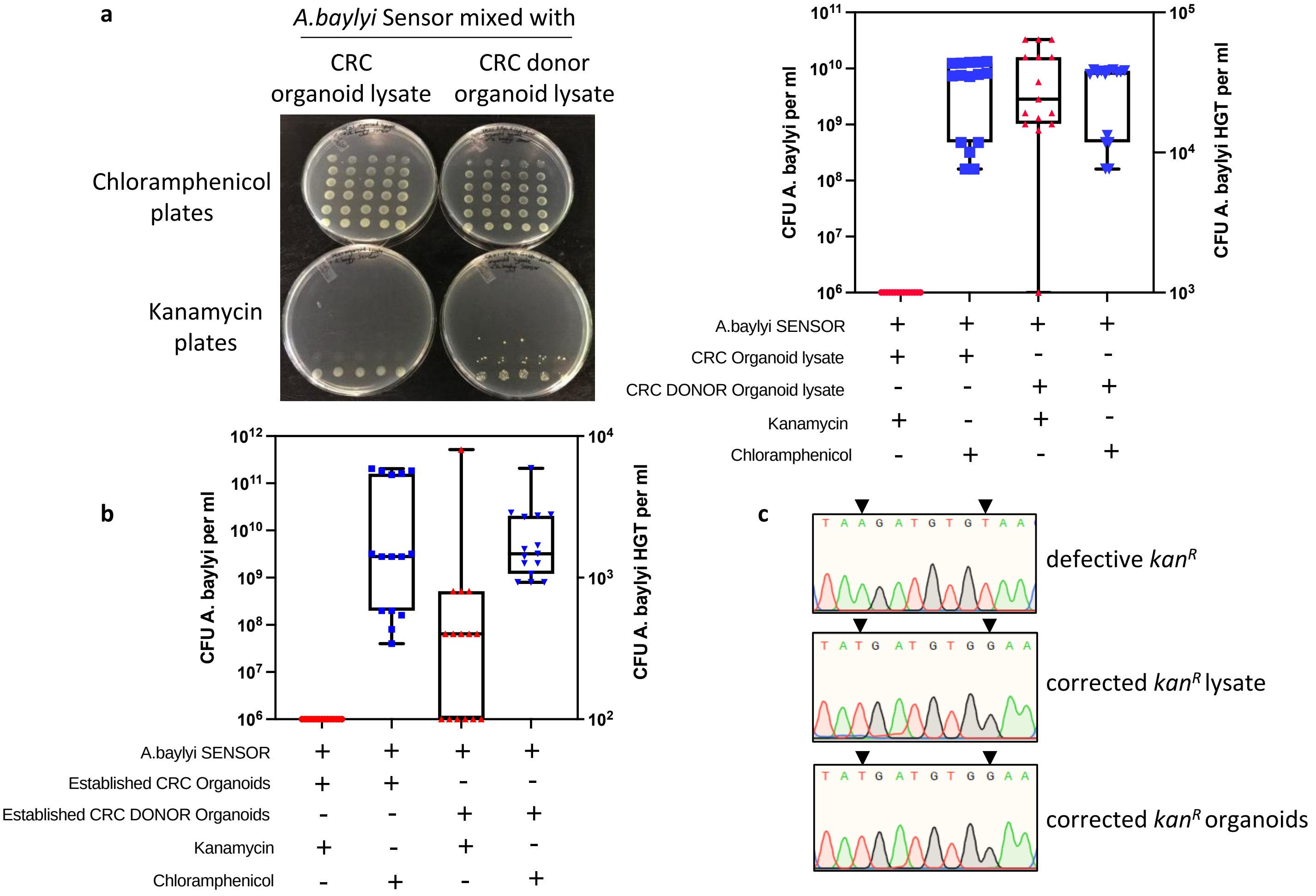


**Extended data figure 3** Sensor detection of donor DNA from BTRZI CRC organoids. *A.baylyi* sensor bacteria are constitutively Chloramphenicol resistant, hence *ChlorR* CFU (CFU *A.baylyi*/ml, in blue) provides a read-out of total *A.baylyi* present. In contrast, Kanamycin resistant sensor bacteria rely on incorporation of donor DNA from CRC organoids to correct the defective *kan* gene and enable growth on kanamycin selection plates (CFU *A.baylyi* HGT/ml, in red). **a** Recombination with lysate from CRC donor organoids enables growth of *A.baylyi* sensor on Kanamycin plates. Shown here with representative plates and CFU analysis. **b** After co-culturing established CRC donor organoids with *A.baylyi* sensor, recombination with donor DNA from CRC donor organoids enables growth of *A.baylyi* sensor on kanamycin plates. **a,b,** Fig3 contains the same data presented here but with average CFU/stool and as HGT rate (CFU *A.baylyi* HGT per ml/CFU *A.baylyi* per ml*)*, n=5 independent experiments each with 5 technical replicates. **c** Representative Sanger sequencing chromatograms of PCR amplicon covering the region of the *kan* gene containing informative SNPs to highlight the difference in sequencing DNA isolated from parental *A.baylyi* sensor bacteria compared to *A.baylyi* colonies isolated from kanamycin plates following mixing with donor organoid lysates or viable organoids.

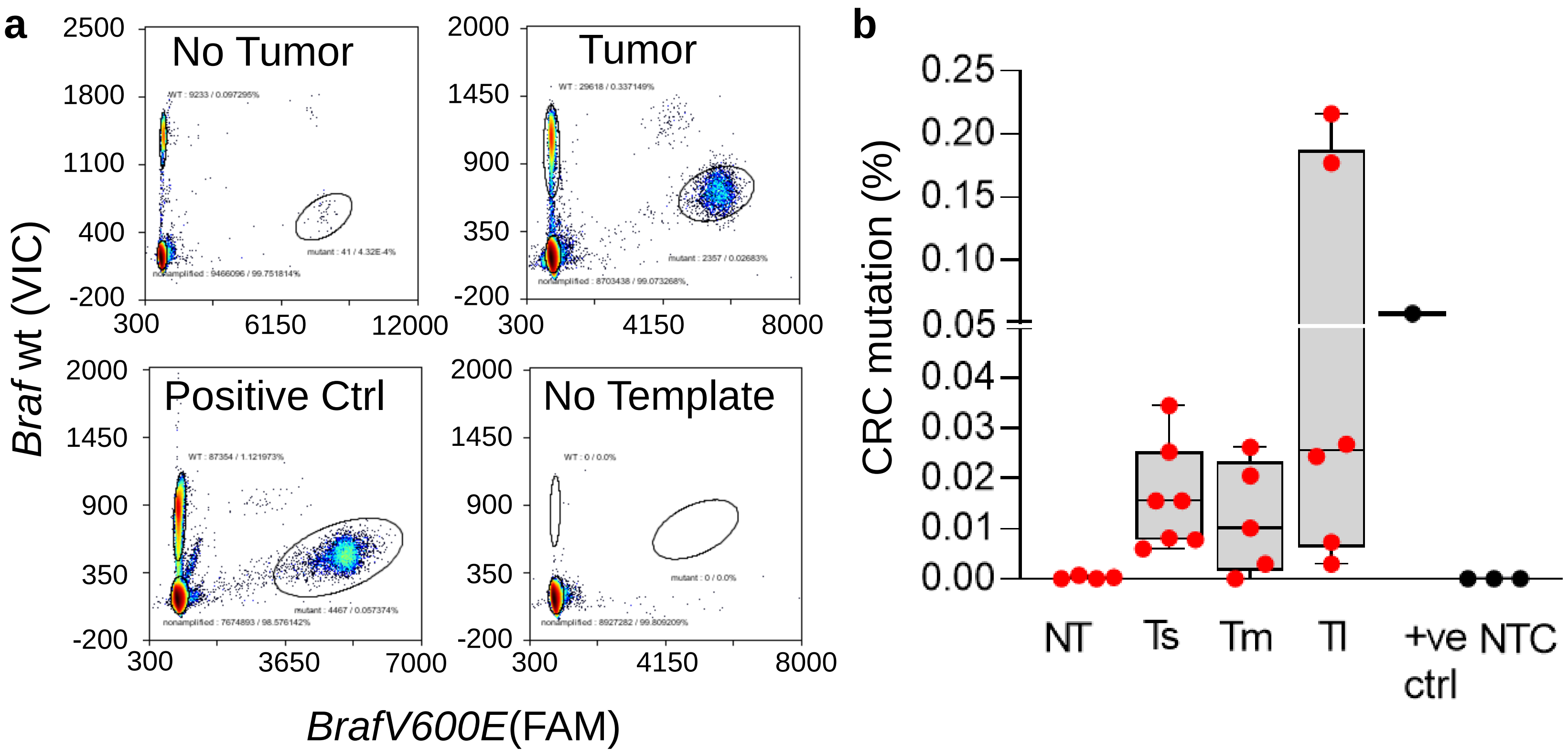


**Extended data figure 4.** High sensitivity digital droplet PCR (ddPCR) detection of CRC mutation (*BrafV600E*) in stool DNA isolated from tumour bearing animals (n=3-4 mice/group). **a,** Representative images of ddPCR data. **b,** CRC mutation (*BrafV600E)* positive droplets as a % of total droplets. Analysis of no template negative control samples and stool DNA samples from non-tumour bearing animals was used to determine the sensitivity threshold of the assay. Positive control samples contain 10% *BrafV600E* gDNA spiked into stool DNA sample from non-tumour bearing animal. NT, no tumour; Ts, small tumour; Tm, medium tumour; Tl, large tumour; NTC, no template PCR negative control.

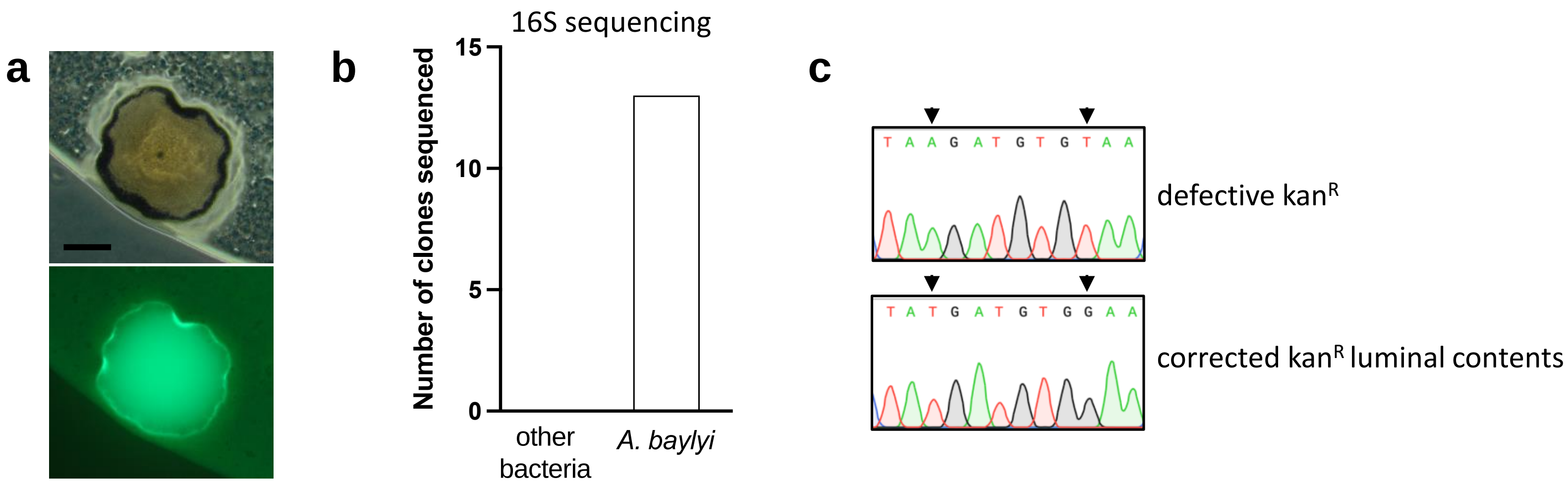


**Extended data figure 5** Horizontal gene transfer is detected in luminal contents from mice bearing BTRZI CRC donor tumors after rectal dosing of *A.baylyi* sensor bacteria. **a** Recombined A.baylyi transformants are GFP positive on kanamycin/chloramphenicol/vancomycin selection plates, scale bar 500 µm. **b** Representative Sanger sequencing chromatograms of PCR amplicon covering the region of the *kan* gene containing informative SNPs to highlight the difference in sequencing DNA isolated from parental *A.baylyi* sensor bacteria compared to *A.baylyi* colonies isolated from kanamycin/chloramphenicol/vancomycin plates in luminal contents from mice bearing BTRZI CRC donor tumors after rectal dosing of *A.baylyi* sensor bacteria

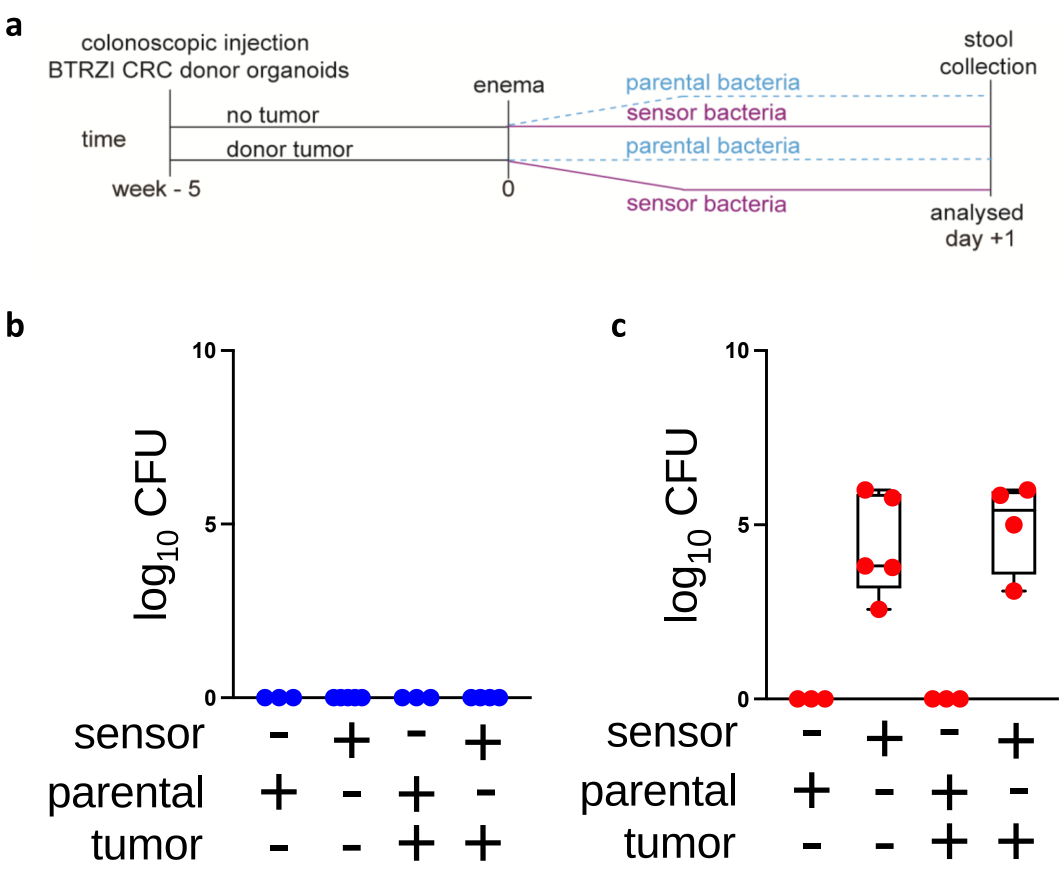


**Extended data figure 6** Horizontal gene transfer is not detected in stool from mice bearing BTRZI CRC donor tumors after rectal dosing of *A.baylyi* sensor bacteria. **a** Schema depicting *in vivo* HGT experiment: generation of BTRZI-*KRAS-kanR* (CRC donor) tumors in mice via colonoscopic injection of CRC donor organoids with tumor pathology validated by H&E histology, administration of parental or sensor *A.baylyi* and stool collection 1 day after enema. **b,c** Rectal delivery of *A.baylyi* sensor to mice bearing CRC tumors results in no detection of HGT in *A.baylyi* sensor bacteria. Data points represent the average CFU per stool from 4 stools per mouse grown on **b** kanamycin/vancomycin selection plates (*A.baylyi* sensor HGT) or **c** chloramphenicol/vancomycin selection plates (total *A.baylyi* sensor), n=3-5 mice/group. Limit of detection 80 CFUs.
